# microbetag: simplifying microbial network interpretation through annotation, enrichment tests and metabolic complementarity analysis

**DOI:** 10.1101/2024.10.01.616208

**Authors:** Haris Zafeiropoulos, Ermis Ioannis Michail Delopoulos, Andi Erega, Aline Schneider, Annelies Geirnaert, John Morris, Karoline Faust

## Abstract

Microbial co-occurrence network inference is often hindered by low accuracy and tool dependency. We introduce microbetag, a comprehensive software ecosystem designed to enhance network annotation. Nodes (taxa) are enriched with phenotypic traits, while edges represent metabolic complementarities, highlighting potential cross-feeding relationships.

microbetag’s online version relies on microbetagDB, a database of 34,608 high-quality genomes with detailed annotations. A stand-alone tool allows users to apply microbetag to custom reference genomes/bins/MAGs. Additionally, MGG, a CytoscapeApp, offers a streamlined, user-friendly interface for network retrieval and visualization. microbetag effectively identified known metabolic interactions and serves as a robust hypothesis-generating tool.

## Background

Most microbial species live only in communities and most natural microbial communities consist of hundreds or even thousands of species. Each species exhibits a unique repertoire of reactions and adapts to various niches, each with specific nutrient and environmental requirements. Depending on the net fitness effects that result for the taxa involved, interactions range from cooperation, competition, parasitism, commensalism and amensalism (1). Microorganisms can interact by competing for or exchanging metabolites. The latter interaction mechanism can involve either one-way (unidirectional) or two-way (bidirectional) exchanges of metabolites.

High-throughput sequencing (HTS) has provided insight into the diversity and composition of microbial communities. Uncultivated species can now be detected, and their features can be inferred from their genomic information (2). Moreover, the composition of thousands of microbiome samples is now accessible allowing for the inference of patterns across large sets of samples. A widely used approach to extract such patterns is the creation of microbial co-occurrence (aka association) networks based on microbial sequencing read data (amplicon and/or shotgun) (3,4). Several approaches are available for co-occurrence network inference based on the assessment of similarity, dissimilarity and correlation (e.g., CoNet (5), SparCC (6)) and conditional dependency identification, which allows reducing the number of indirect edges (e.g., SpiecEasi (7), FlashWeave (8)). Nevertheless, microbial network inference encounters various challenges (9,10). Each inference approach comes with its own assumptions and parameter settings, leading to variations in network structure. The choice of the association measure and data preprocessing techniques as well as the handling of sparsity and zero inflation all influence the resulting network. As a consequence, the result of network construction is tool-dependent (11,12). Moreover, microbial network inference inherits the challenges of sequencing data and analysis (e.g., sampling scale, compositionality, tuning of parameters linked to sequencing data processing) and the returned network is often a “hairball” of densely interconnected taxa. Thus, additional analysis is necessary to generate testable hypotheses (10).

The comparison of interactions predicted by microbial networks with a collection of known interactions has underscored their low accuracy for this task (13). Data integration has been suggested to help interpret edges in microbial networks (10). In addition, clusters in microbial networks have been demonstrated to detect key drivers of community composition (14) and several algorithms and implementations are available to identify them (15). However, data integration approaches available for microbial networks are so far limited.

Metabolic networks are comprehensive representations, in a mathematical form, of biochemical reactions occurring within an organism (16–18). Over the last decade, (semi-) automated approaches support the fast generation of such reconstructions (e.g. (19,20)). A metabolic network’s “seed set” is the set of compounds that, based on the network topology, cannot be produced by the organism and needs to be acquired exogenously (21). Such seeds might be independent, i.e. they cannot be produced by any other biochemical reaction in the metabolic network, or they can be interdependent forming groups of seed compounds. Seeds are a useful proxy for the essential nutrients of an organism (21,22). Based on the seed concept, several graph theory-based metrics (indices) have been described to predict species interactions directly from their metabolic networks’ topologies (23–26).

Metabolic complementarity among species reflects the potential for cooperation through cross-feeding. In contrast, metabolic competition refers to the metabolic overlap between two species leading to exploitative competition, e.g. for nutrients. Seed and non-seed compound sets can be used to compute complementarity and overlap indices. The examination of such indices can indicate metabolic interactions that may drive the patterns observed in co-occurrence networks.

To explore whether a species may benefit from a partner, it is helpful to move from the network to the pathway level and check whether their pathways complement each other. Here, we rely on a naive approach exporting all possible complements between a pair of species based on their KEGG ORTHOLOGY (KOs) annotations and the KEGG MODULES database (27).

Here, we present microbetag, a microbial network annotator and software ecosystem that exploits several sources of information to enhance or reduce the confidence in the associations suggested by the network, thereby generating hypotheses for further investigation both at the taxon pair and the community level. microbetag serves as a comprehensive platform that provides information on taxa along with their potential metabolic interactions from multiple channels (see “Methods”). The key concept here is the reverse ecology approach (28). Reverse ecology leverages genomics to explore community ecology with no *a priori* assumptions about the taxa involved. The reverse ecology framework enables the prediction of ecological traits for less-understood microorganisms and their interactions with others (29). microbetag annotates a user’s co-occurrence network by integrating phenotypic traits of the taxa present in the network (nodes) and by mapping potential metabolic interactions onto their associations (edges). microbetag is accompanied by a Graphical User Interface (GUI) implemented as a Cytoscape (30) App (31) providing a user-friendly environment to investigate annotations in a straightforward way. Its online version depends on precalculated annotations for more than 35,000 high quality reference genomes stored in the microbetagDB. All annotations present in microbetagDB are also available through an Application Programming Interface (API). microbetag’s source code is distributed under a GNU GPL v3 license and available on GitHub. Documentation and further support on how to use microbetag is available at the <u>documentation website</u>. To the best of our knowledge there is no software with which microbetag could be compared to directly. To validate our annotations, we used a recently published network with partially known interactions (32). We then present the main features of its interface and discuss an application example.

## Results

### Overview of the microbetag software ecosystem

The microbetag software ecosystem consists of five main modules: a) microbetagDB consisting of the precalculations for 34,608 reference genomes and their pairwise combinations (see Methods), b) a web server hosting microbetagDB and the microbetag app to annotate a co-occurrence network on-the-fly or to retrieve annotations through an API, c) a Cytoscape app called MGG that enables users to easily invoke the workflow and investigate the annotated network, d) a preprocessing workflow for data sets with more than 1,000 sequence identifiers (OTUs/ASVs/bins etc.) and e) a stand-alone version of the microbetag precalculation steps combined with the microbetag workflow that supports annotating a network based on the user’s reference genomes.

Figure 1 shows a simplified overview of microbetag’s on-the-fly annotation approach. Through the MGG Cytoscape App we provide, the user may load an abundance table and optionally its corresponding network and send a *job* to the microbetag server. If not provided, microbetag will then infer a network using FlashWeave, and it will annotate its nodes (taxa) with phenotypic traits and its edges, which represent significant co-occurrences or mutual exclusions, with potential metabolic complementarities. The annotated network is then returned to the user as a response to their job-query and will be automatically loaded on Cytoscape’s main window. The user can then investigate its annotations further thanks to the MGG features (see “Annotating networks with microbetag” paragraph).

**Fig. 1.**
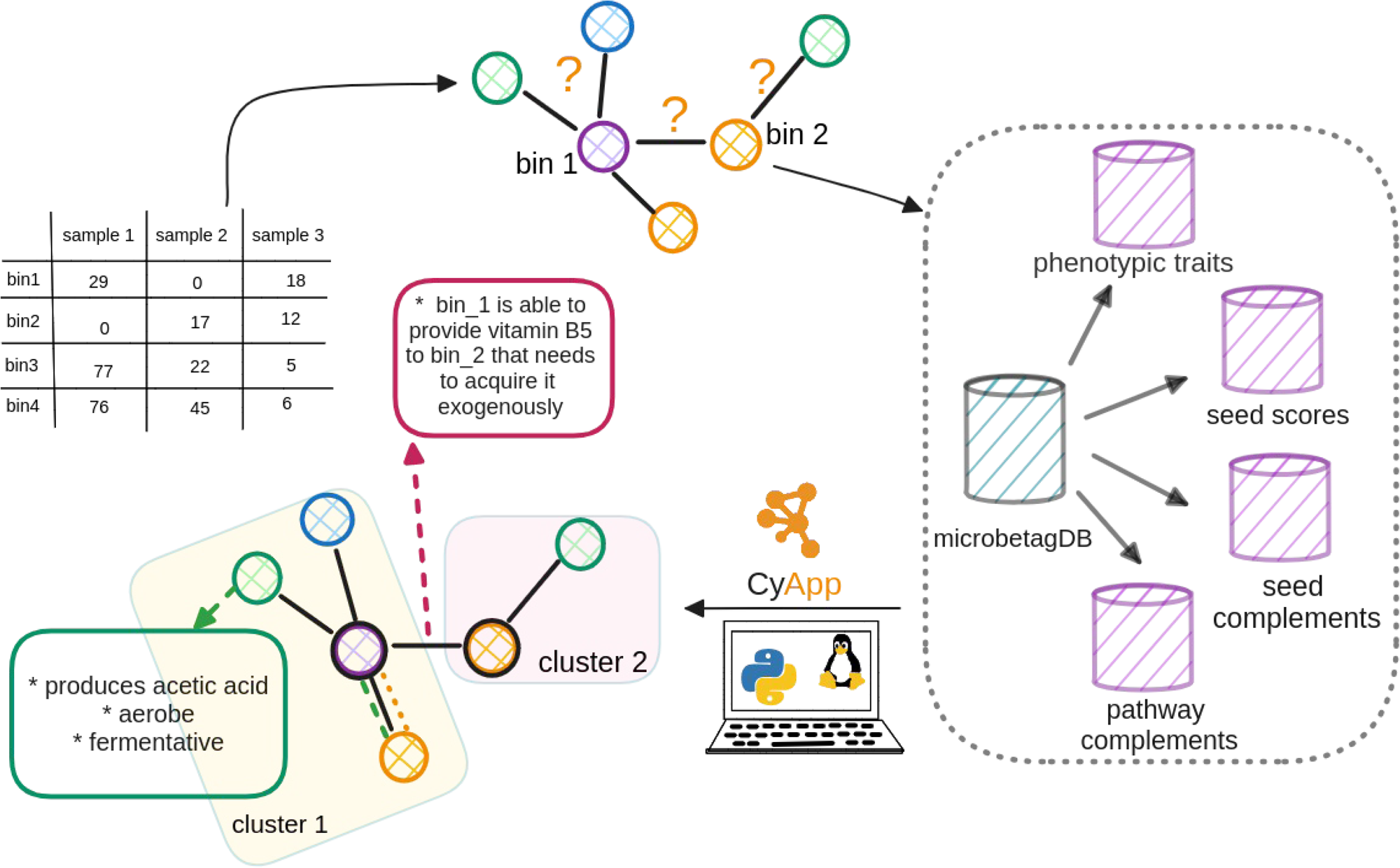
Simplified overview of the microbetagDB-dependent version of microbetag.

### The microbetagDB resource

Currently, microbetagDB includes more than 34,000 genomes (Table 1) along with their corresponding annotations. Most of these genomes represent bacterial taxa (364 taxa are Archaea). The presence/absence of more than 30 phenotypic traits have been predicted for those genomes. About 1.4 billion potential metabolic interactions leading to pathway or seed complementarities have been precomputed as well. There is one order of magnitude more seed complements than those corresponding to pathway complementarities since for all metabolic networks present in microbetagDB all pairwise complements were calculated (33,755^2^) and stored, even if empty. In the case of pathway complementarities, a genome pair is present in the database and thus counted only if a potential complementarity was found. Yet, in the first case, the number of genomes with no potential seed complements ranges from zero to a few dozen. All annotations can be accessed directly from microbetagDB through the API. Seed complements can be interpreted as compounds that are essential for the beneficiary since, by definition, they cannot be produced by its metabolic network, and which can be potentially cross-fed from the donor to the beneficiary.

**Table 1:**
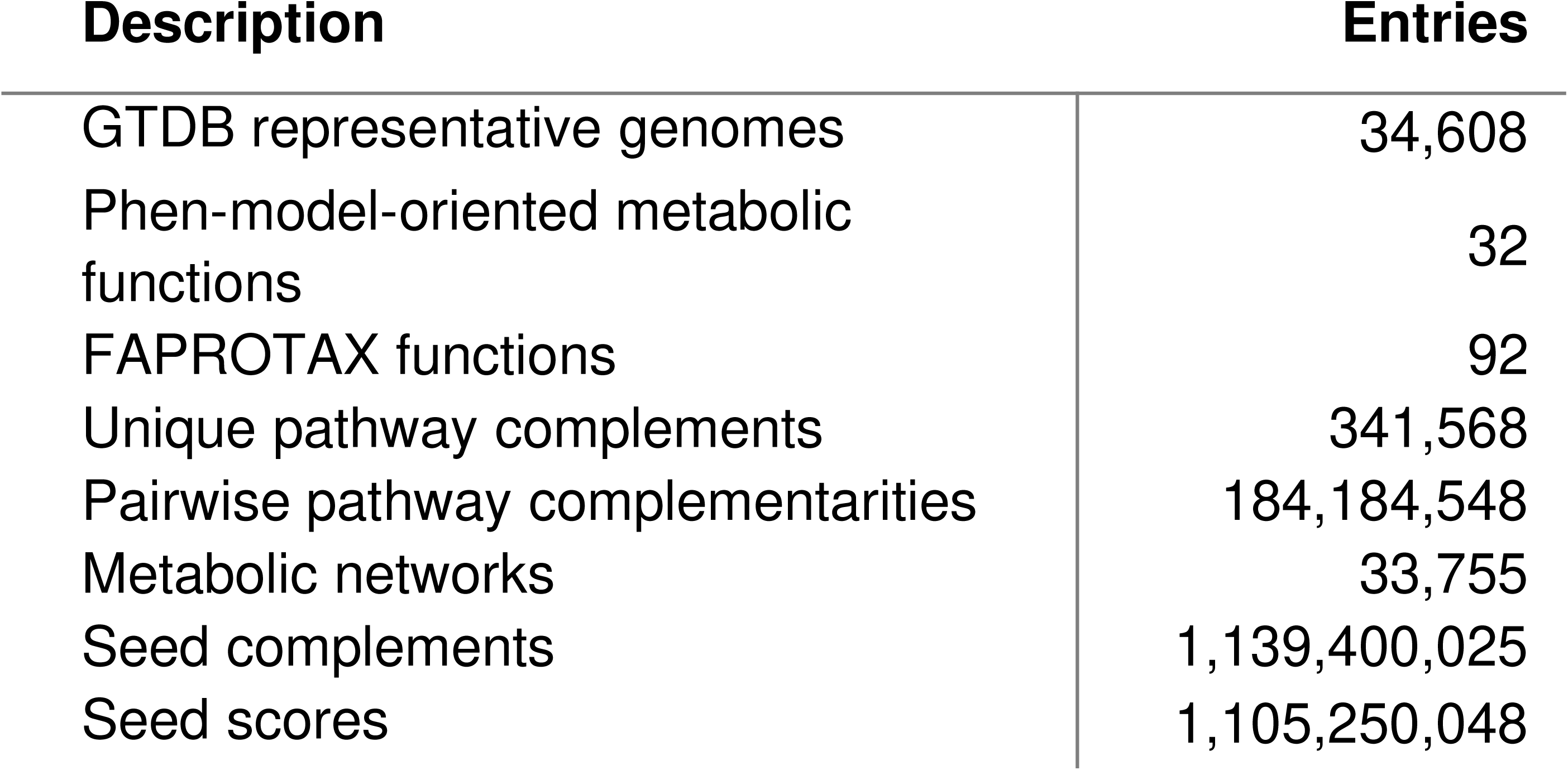
Summary of the data in microbetagDB.

| Description | Entries |
| --- | --- |
| GTDB representative genomes | 34,608 |
| Phen-model-oriented metabolic functions | 32 |
| FAPROTAX functions | 92 |
| Unique pathway complements | 341,568 |
| Pairwise pathway complementarities | 184,184,548 |
| Metabolic networks | 33,755 |
| Seed complements | 1,139,400,025 |
| Seed scores | 1,105,250,048 |

### Annotating networks with microbetag

Thanks to MGG, a Cytoscape App specifically developed to streamline the use of microbetag, running microbetag is straightforward and requires no bioinformatics skills. In the simplest case, microbetag only takes an abundance table with taxonomic assignments as input. When the taxonomy scheme being used is not one among the GTDB, Silva or the GTDB-oriented taxonomy for 16S rRNA amplicon data (see “microbetag preprocessing”), the most time-consuming step of the workflow is the one mapping the user’s taxonomy to an NCBI Taxonomy Id and from that to GTDB representative genomes. Network inference can be a computationally intensive step too, particularly as the number of taxa in the abundance table increases. To enable annotation of large data sets, a stand-alone preprocessing workflow is provided with microbetag. The user can either assign their amplicon data to the GTDB-oriented taxonomy and/or reconstruct a network locally. Once a network is available and the taxonomy provided is among the standard ones for microbetag, the computational time required for annotation ranges from several seconds to a few minutes based on the user’s settings. An annotated network in .cx2 format is returned, which can be viewed in Cytoscape.

To enable the microbetag approach with custom/local genomes, instead of mapping user’s taxa to the GTDB representative ones, a microbetagDB-free version is also available. In this case, the user has to pull a Docker image of microbetag that supports all the annotation steps for user-provided genomes. microbetag will first perform all the genome annotation steps and the metabolic model reconstruction if needed depending on the user’s settings. For example, in case a user has already annotated their genomes with KEGG, or they have already reconstructed their corresponding metabolic networks, the corresponding precalculation steps may be omitted. Then, the microbetag workflow (see “Methods”) will be performed to return an annotated network. The computing resources for such a task can be quite extensive (see “Running times”). Once completed, the annotated network will be saved as a .cx2 file and can be visualized in Cytoscape using the MGG app.

Tutorials for the different scenarios for which one can use microbetag, frequently asked questions and hints to address the idiosyncrasies of various data sets are available on the documentation web site (https://hariszaf.github.io/microbetag/) while a Matrix community (https://matrix.to/#/#microbetagcommunity:matrix.org) allows users to exchange experiences and ask for more specific help. In the following two sections, we present a validation and use case, highlighting our approach’s potential.

MGG allows the user to import data, retrieve an annotated network and investigate the annotations through a series of CyPanels both for node and edge annotations. Figure 2 shows an example of the CyPanels. In the node panel (Fig. 2.A), the node name, taxonomy, NCBI Taxonomy Id and GTDB genome to which the sequence was mapped can be viewed. Depending on the user’s settings and the available annotations for a node, genome-or literature-based predictions may be present. Further, the trait groups mentioned in “Groups of phenotypic traits” are on top of this panel allowing for the selection of the nodes carrying either one among several attributes (OR logical relationship) or all of them (AND). Accordingly, in the edge panel (Fig. 2.B), the set of beneficiary taxa is specified along with their corresponding GTDB representative sequence identifiers. Pathway and seed complementarities are shown each in a table. Potential metabolic interactions are shown in a sub-table entitled with the genome pair under consideration, as several GTDB genomes may have been assigned to a node. In case of pathway complementarities, these tables consist of six columns: a) the KEGG MODULE id of the module to be completed, b) its description, c) a more general metabolic category to which the module belongs, d) the complement itself as a list of KEGG terms, e) the alternative that now represents a complete module in the beneficiary and f) a URL that points to a colored KEGG map highlighting the complement. If clicked, the user’s default browser pops up showing a colored KEGG map as shown in an example in Figure 2.C.

Last, MGG allows testing for enrichment or depletion of the phenotypic traits assigned to the nodes in each of the network clusters. Clusters are returned from manta (15) while performing the microbetag workflow if requested or users may assign them manually or using another network clustering algorithm. For thorough instructions on how to use MGG and microbetag the reader may visit the documentation web site (https://hariszaf.github.io/microbetag/docs/cytoApp/).

**Fig. 2.**
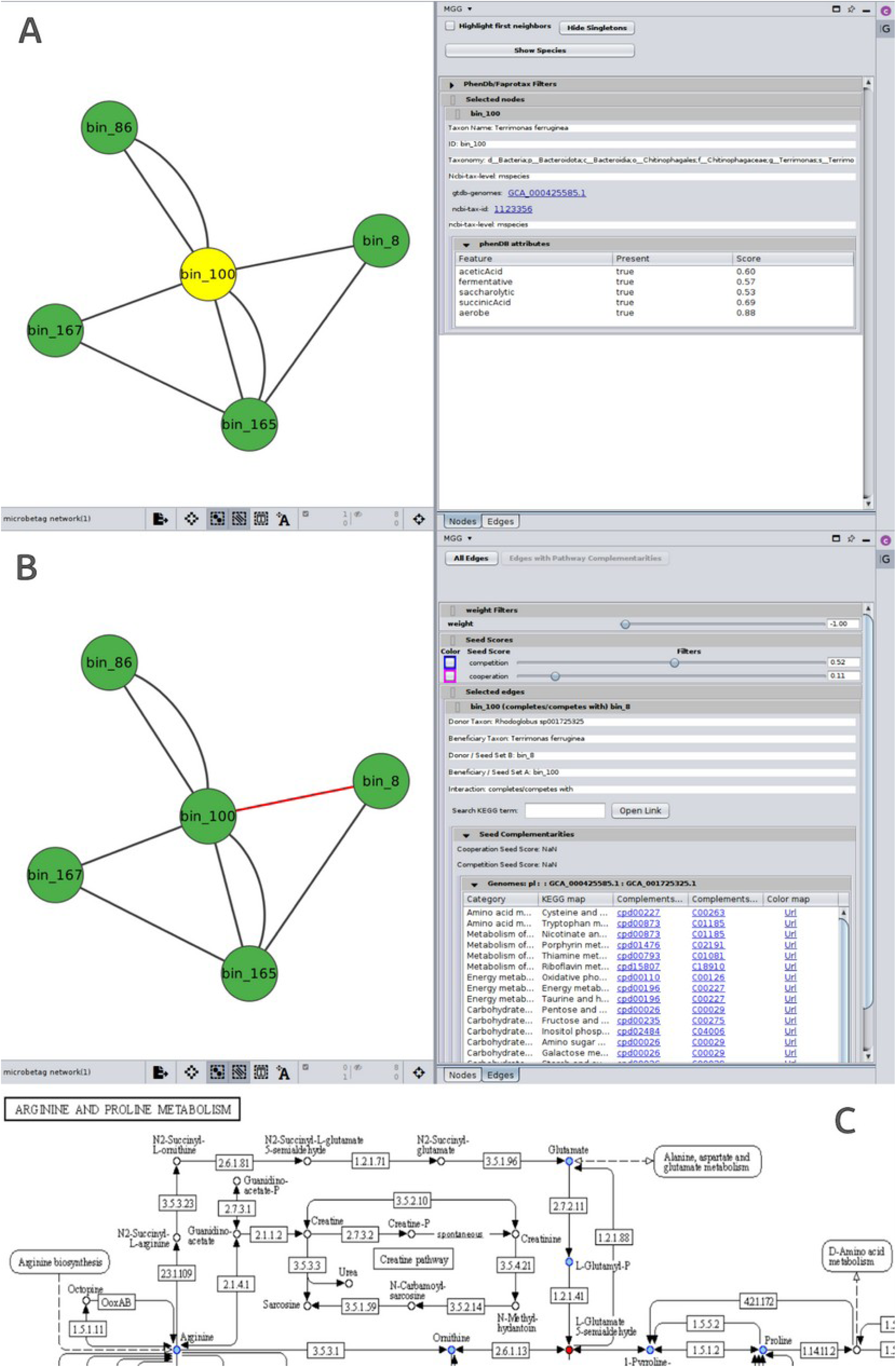
CyPanels of the MGG Cytoscape app. **A** *Nodes* panel displays the annotations of each taxon (node) mapped to one or more GTDB genomes. In this example, genome-based predicted phenotypic traits are shown along with their prediction score. **B** *Edges* panel displays the list of potential metabolic complementarities between two nodes, specifying which is the potential donor and the potential beneficiary taxon, thus giving a direction to the corresponding edge in the graph. There are two cases of complementarities in microbetag. The first are seed complementarities, which are computed based on the species’ reconstructed metabolic networks (case shown above). *Seed complementarities* are first exported based on ModelSEED complements (column three) and mapped to KEGG COMPOUNDS (column four). The provided URL points to a colored KEGG map that highlights seeds complementing a beneficiary’s metabolic network. The second are pathway complementarities. These are directly computed from the KEGG annotated genomes and do not involve ModelSEED ids since the metabolic network construction step is not necessary in this case. In brief, seed complementarities represent essential compounds that the beneficiary cannot produce, and pathway complementarities represent enzymes that fill gaps in a beneficiary’s pathway. **C.** Part of a colored KEGG map returned based on the seed complementarities. Compounds that belong to the beneficiary taxon are colored in cyan while the potential complementing seeds from the donor are highlighted in red.

### Validation of microbetag on a network with known interactions

We validated microbetag using the correlation network of Hessler et *al.* (32), which describes mine tailing-derived laboratory microbial consortia. In this study, *Variovorax*, a thiamine producer, and its co-occurrence with a series of thiamine auxotrophs are discussed. The authors tested network predictions by performing co-culture experiments measuring the thiamine production. Sequence bins corresponding to network nodes and the original network were obtained from the authors. Using GTDB-tk (33), bins were assigned GTDB taxonomies; those retrieved were added to the original network, which was then annotated with microbetag. Supplementary Fig. B2 highlights bin_55 that corresponds to *Variovorax* as well as its first neighbors. The annotated network is available on microbetag’s GitHub repository. GTDB-tk returned GCA_001899795.1 as the one closest to bin_55, assigning it to *Variovorax* sp001899795. microbetag then suggested that this specific genome corresponds to an aerobe (34) that can grow autotrophically (35) and consume D-glucose, while producing ethanol and lactic acid (36). Last, the Type VI secretion system was suggested to be available on its genome (37).

Hessler et *al.* (32) argue that *Variovorax* is an important thiamine source and can supply neighboring species that cannot produce it (auxotrophs). Indeed, microbetag suggested several thiamine-related potential seed complements among the potential metabolic interactions between *Variovorax* and its neighbors (Table 2). Potential interactions were also found in some cases between the neighbors themselves (Supplementary Table 2). The authors also argue that isolates of that *Variovorax* strain required the addition of pantothenic acid to grow. However, based on the KEGG annotation of the genome that bin 55 was mapped to, it contains both KEGG MODULES related to pantothenate biosynthesis, M00119 (valine/L-aspartate ⇒ pantothenate) and M00913 (2-oxoisovalerate/spermine ⇒ pantothenate) whereas other genomes are not capable of either one or any of those reactions (Supplementary Table 2). This example highlights a challenge of the microbetag approach. The complementarities between the nodes would have been different if bin 55 was classified as another *Variovorax* genome, with incomplete modules. For example, if GCF_001577265.1 was picked and its complementarities with the neighboring taxa were retrieved, it would have revealed that all its neighboring species can provide it with pantothenate (C00864) as suggested by their seed complementarities (see colored map).

**Table 2:**
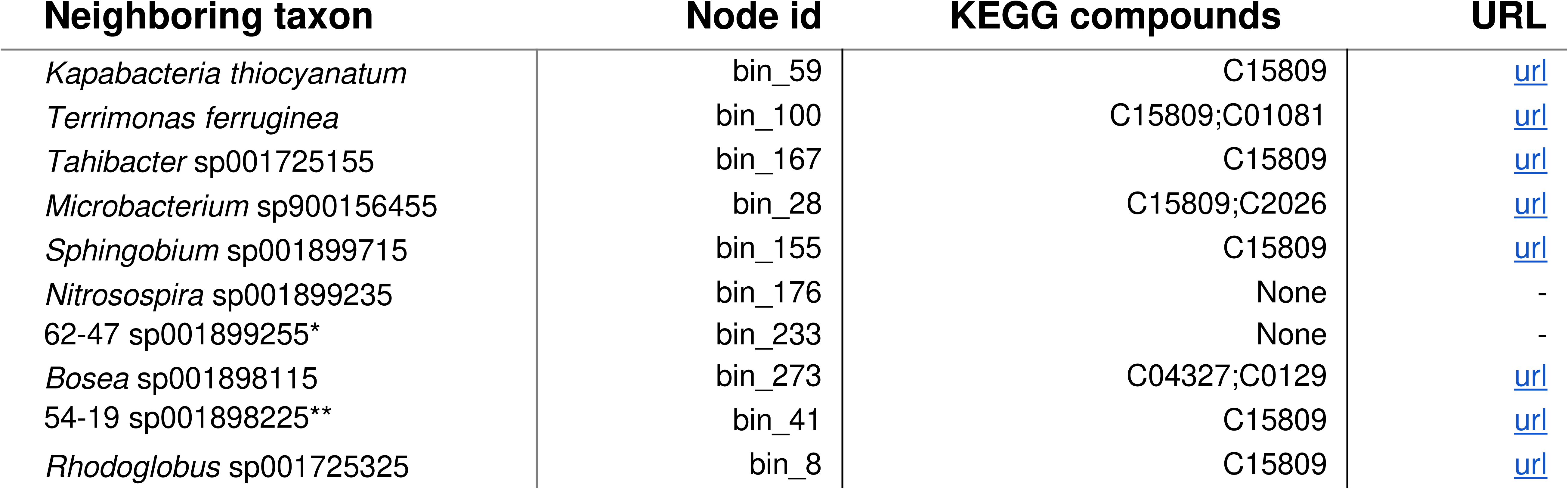
Thiamine biosynthesis related seed complements between *Variovorax* and its first neighbors in the network of Hessler *et al.* (32). In all cases but the one with *Bosea, Variovorax* is the potential donor of seeds for the rest of the community members. KEGG compound ids correspond to the following compound names C15809: Iminoglycine; C01081: Thiamin monophosphate; C20246: 2-[(2R,5Z)-2-Carboxy-4-methylthiazol-5(2H)-ylidene]ethyl phosphate; C04327: 4-Methyl-5-(2-phospho-oxyethyl) thiazole.

### Application of microbetag to in-vitro data of infant gut microbiota

In their study, Cabrera et *al.* (38) showed that iron supplementation combined with either inulin or short-chain galacto-oligo oligosaccharides (scGOS) and long chain fructo-oligosaccharides (lcFOS) leads to a higher relative abundance of bifidobacteria, increased production of acetate, propionate, and butyrate, and a significant shift in microbial composition compared to non-supplemented microbiota. In this way, enteropathogens that are usually increased due to iron supplementation, a common approach to prevent anemia, were reduced.

To analyze these data further, we first ran the microbetag preparation step to map the ASVs in the 16S rRNA sequencing data of this study to GTDB-based taxonomic assignments. Then we inferred the network with FlashWeave (wrapped by microbetag) and annotated it using microbetag. Figure 3A shows the microbetag-annotated co-occurrence network for microbiota from donor 3, an eight-month-old female infant, whose fecal microbiota was used to inoculate the continuous *in vitro* experiment. After a stabilization period of 7 to 13 days, iron and scGOS/lcFOS were continuously supplied for 6 to 9 days, and this treatment set-up was repeated twice. microbetag predicted several taxa to be butyrate producers. Interestingly, most of the associations involving butyrate producers are negative, likely reflecting that they represent a different stage of the succession. At the same time, a single butyrate producer (*Flavonifractor plautii*; ASV0012) had several positive associations. Microbetag reports that *F. plautii* and its neighbor species *Eggerthella lenta* (ASV0107) have several pathway and seed complementarities (Fig. 3.B), in particular for Coenzyme A biosynthesis, which is involved in the bacterial butyrate production pathway (39). More specifically, *F. plautii* requires alanine and pantothenate to be exogenously acquired, while *E. lenta* is capable of producing both of them. Thus, the analysis with microbetag results in a testable hypothesis: the observed increase in butyrate production and *Flavonifractor* in infant microbial communities upon FOS treatment *in vitro* (see Figures 5B and 7 in (38)) may be, at least to some extent, due to cross-feeding partners boosted by the treatment that complement metabolic pathways involved in butyrate production (Fig. 3.C, 3.D). Relevant files for the use case can be found at microbetag’s main GitHub repository.

**Fig. 3.**
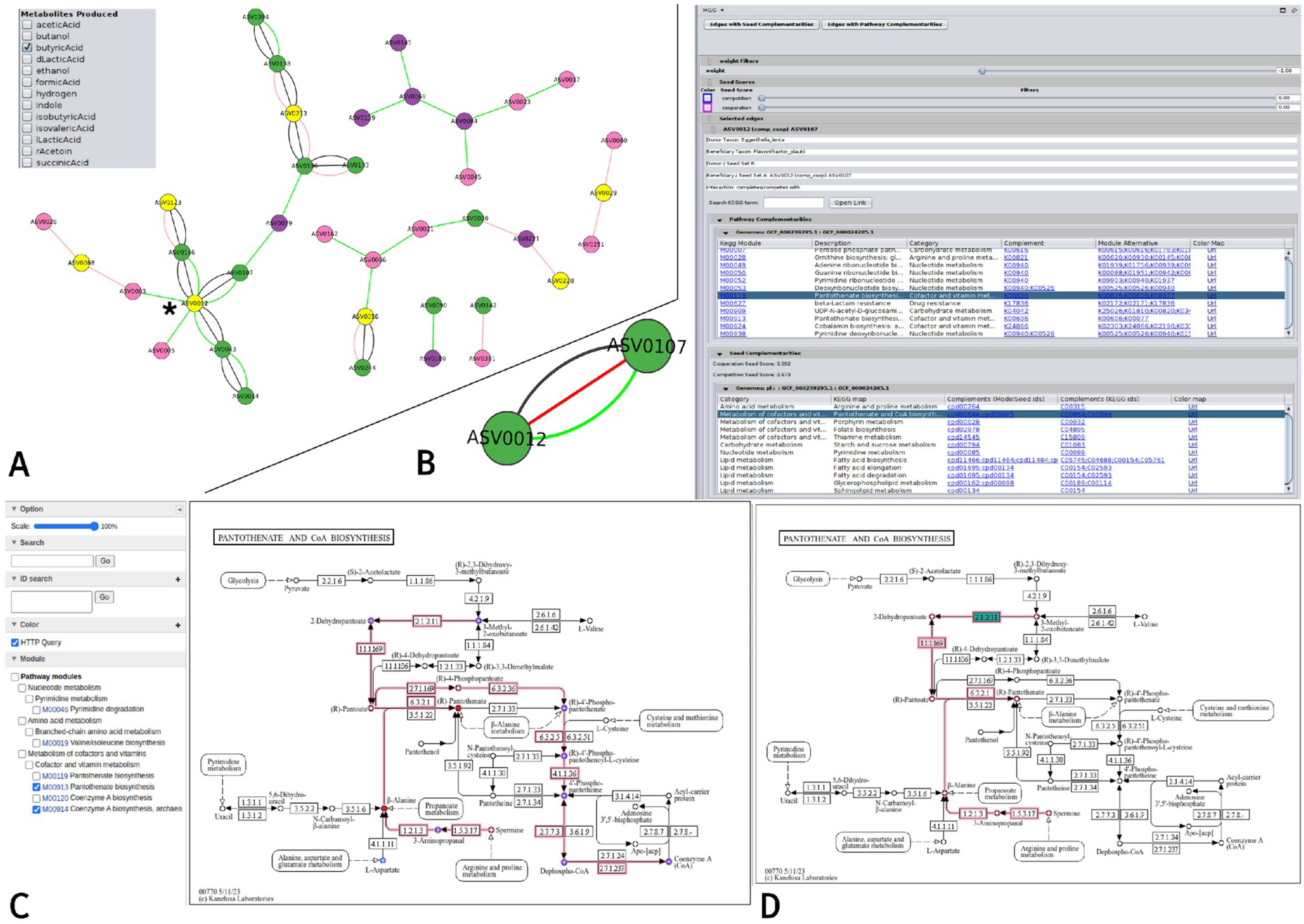
microbetag analysis of a dataset of infant gut microbiota studied *in vitro* (38). **A.** The microbetag-annotated co-occurrence network as viewed in Cytoscape with the MGG visual style. Taxa predicted to be butyrate producers are highlighted in yellow. The asterisk (*) points to *Flavonifractor plautii*. **B.** microbetag edge annotations considering *F. plautii* (ASV00012) as the potential beneficiary and *Eggerthella lenta* (ASV0107) as a potential donor. In this case, both pathway and seed complementarities are available and complements related to pantothenate (vitamin B5) biosynthesis, and thus Coenzyme A biosynthesis are present. **C.** Colored Pantothenate and CoA biosynthesis KEGG map returned by microbetag highlighting potential seed complementarities provided by *E. lenta*; the two seeds of *F. plautii* that *E. lenta* may provide, red β-alanine and (R)-pantothenate, are highlighted in red. **D.** The same KEGG map suggests a missing enzyme (2.1.2.11) of *F. plautii* for producing pantoate that could be acquired from *E. lenta* and that would enable *F. plautii* to produce pantothenate from pantoate (pathway complementarity).

### Pathway complementarity statistics

In its current version (v1.0.1) microbetag supports complementarities for 491 KEGG modules. These modules have been parsed to identify 23,592 alternatives, i.e. sets of KO terms that ensure the module to be complete (see “Pathway Complementarity” paragraph in “Methods”). Using the 16,902 GTDB genomes with a .faa file, a total of 184.2M pairwise pathway complementarities were retrieved, where a complementarity or complement is a combination of KOs that complete an alternative and thus enable a KEGG module. Theoretically, one would expect 285.5M pairwise complementarities, yet not every possible pair of genomes has a complement. More specifically, 237,075 unique complements, i.e., combinations of KOs that complete an alternative and thus enable the module, were found to complete 6,467 alternatives in a total of 341,568 unique pathway complementarities. The latter is due to the fact that the same complement may complete more than one module at the same time. It is worth mentioning that almost 1/7 of those are related to the Embden-Meyerhof pathway of glycolysis. This is expected since 13,440 of the alternatives are related to this module. However, only 2,921 of those alternatives had potential complements based on our genomes. However, for the vast majority of the modules, all the enumerated alternatives were found to be potentially completed by another taxon (Fig. 4.A).

In addition, 56 modules were found to carry more than 10 alternatives, covering the 96% of the alternatives observed (Fig. 4.B, 4.C). Yet, 49,944 of the total 341,568 unique complementarities come from the remaining 435 modules. We also filtered the unique complements for those requiring more than 4 KOs from the donor species; we assume that the more KO terms need to be shared, the more challenging it is for the complementarity to actually occur. We found that out of the 215,883 cases requiring more than 4 KOs, the 185,937 represent cases of the 56 modules with the highest number of alternatives.

The KEGG modules’ definitions are global for all domains of life. However, some *KO terms* are only present in Animals or Plants etc. That explains partially the alternatives that are never completed. For example, in case of md:M00002 the term K12406 is only present in Animals (https://www.genome.jp/kegg-bin/show_brite?htext=br08611&pruning=join_brite&brel=off&hier=20&highlight=join_brite&mapper=taxmap-s-p%20K12406). Thus, from the total 24 alternatives for that module, only 12 are present in microbetagDB, as expected. In addition, there are also *KO modules* that can occur only in Plants and/or Eukaryotes. That explains to a great extent why for 168 modules, no alternatives were completed at all. For example, module M00372, the “Abscisic acid biosynthesis”, (https://www.genome.jp/kegg-bin/show_brite?htext=br08611&pruning=join_brite&brel=off&hier=20&highlight=join_brite&mapper=taxmap-s-p%20M00372) is not present in microbetagDB as expected. There are also 3 KEGG modules with more than 1000 alternatives.

**Fig. 4.**
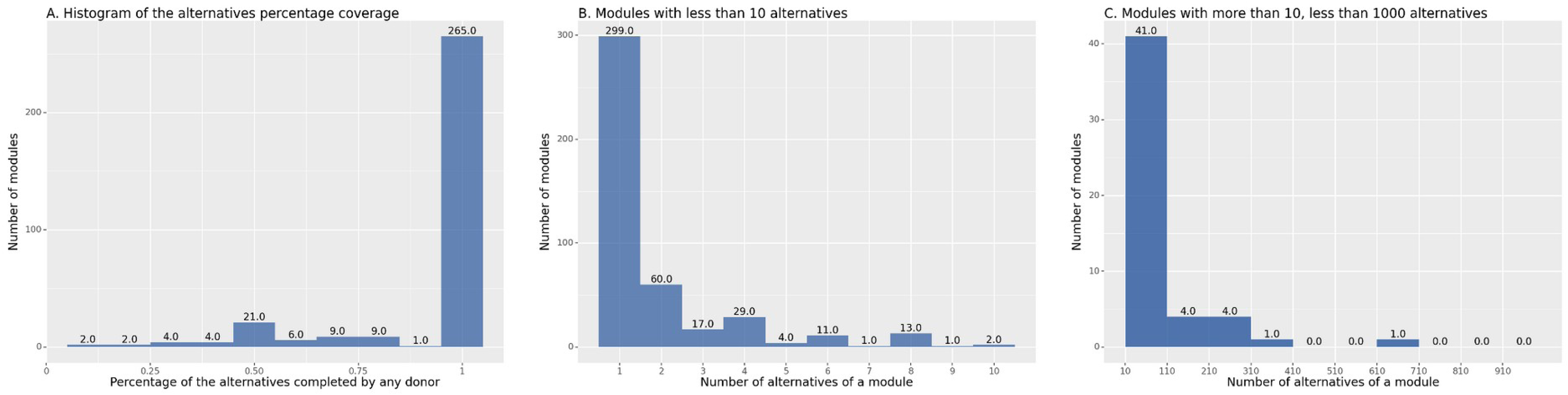
Distribution of the percentage of potentially completed alternatives for the KEGG modules available on microbetagDB and of the number of their total alternatives. **A.** For the vast majority of the modules all their different alternatives can be completed by a potential donor species (265). A sum of 323 modules out of the 491 available on KEGG do have at least one way to complete an alternative in the microbetagDB framework. **B.** and **C.** Most of the KEGG modules have 1 or 2 alternatives but there is a set of 132 modules for which the number of their alternatives range from 3 to 13,440 (Embden-Meyerhof pathway, M00001).

## Running times

### On-the-fly

Using the abundance table from the validation case (vitAbund.tsv, 120 bins) annotated with the GTDB taxonomy scheme and its corresponding network file (edgelist.tsv, 102 nodes, 251 edges) it took 40’’ for microbetag to return an annotated network; microbetag was able to annotate 55 of its nodes and 93 of its edges. Next, using a different abundance table containing 16 samples and 78 Silva-annotated ASVs (testAbundMeta.tsv), microbetag took 67’’ to infer a co-occurrence network with 23 nodes and 14 edges using the FlashWeave sensitive approach. It managed to annotate 7 of the nodes and 1 of the edges in this network. We then ran the same analysis with an abundance table that had the same ASVs as in the previous example but with 84 samples (testAbund.tsv). This time, it took 74’’ for microbetag to return a co-occurrence network of 78 nodes of which 19 got annotated and 118 edges of which 5 were annotated. We then used the returned network and built a 3-columns file for its co-occurrence weight and used it as input and setting the taxonomy scheme to “Other”, microbetag returned an annotated network with 16 annotated nodes and 3 annotated edges.

### Preprocessing tool

We used the preprocessing image with an abundance file of 16 samples and 997 ASVs and annotated them with GTDB-based taxonomy. We also performed the FlashWeave step using the sensitive approach. Using a personal computer of 15 cpus, it took 3’ 52’’ real time while the user time was 24’ 19’’ implying a degree of parallelization. On average 6 cpus were used. Using the taxonomy annotated abundance table directly on the web-app version of microbetag, it returned a network of 240 nodes, 105 of which annotated and 219 edges, 49 of which got annotated in 94’’. When we used both the GTDB annotated table and the network returned from the preprocessing step, it took 40’’. One can exploit the parallel performance of FlashWeave thanks to Julia implementation; in our running no parallelization was used.

### Stand-alone version

Using custom genomes to run microbetag with, requires an important amount of computing resources, especially in two of the performed steps: gene recognition and the KEGG annotation of the genomes. In case ModelSEEDpy is being used for genome-scale reconstruction, then RAST annotation would also be required . In our demo case, with only 7 bins to annotate, it took almost 1 hour for ORFs (running prodigal through DiTing) and about 2 hours for the KOs assignment using a personal computer with 2 CPUs. Reconstructing the metabolic models with carveme is faster and performs in a more robust way since there is no need for establishing any connection with external servers as in the ModelSEED case, where a connection with the RAST server is required.

## Discussion

### Potential and limitations

The previous paragraphs illustrate the potential of microbetag in the interpretation of co-occurrence networks and how it can be used to generate new hypotheses derived from those. However, microbetag benefits the microbiome community in several other ways. The microbetagDB provides a vast number of annotations; 31 predicted traits for more than 30,000 genomes, their metabolic networks along with their corresponding seed sets, potential metabolic complementarities and cooperation/competition scores. Such a resource may support a range of studies; from a more theoretical perspective regarding the distribution of the complements among taxonomic groups or how often a complement potentially appears, to applications such as eco-evolutionary studies and the investigation of interactions. For example, the “Pentose phosphate pathway” module (md:M00004) was the one with the most alternatives [40] for which all alternatives were found to be possible to complete from another species. At the same time, “De novo purine biosynthesis, PRPP + glutamine => IMP” (md:M00048) is the one with the lowest percentage of alternatives completed by any donor (0.06%). A closer look to the latter shows that as part of the definition of the module, there are KOs that can be found only in non Bacteria taxa; e.g. among all the bacterial KEGG genomes, K11787 (phosphoribosylamine--glycine ligase) is present only in *Defluviicoccus* sp. SSA4 while K01587 (phosphoribosylaminoimidazole carboxylase) only in *Rhodoplanes* sp. Z2-YC6860 and *Berkelbacteria* bacterium GW2011_GWE1_39_12). In addition, K11787 is responsible for 3 steps of the module in case of *<u>H. sapiens</u>*, suggesting a significantly different way of implementing the same task/module compared to taxa of lower complexity. This implies an evolutionary spectrum that could be used to further investigate what is going on within prokaryotes too.

For studies with a short number of sequence identifiers the on-the-fly version of microbetag returned annotated networks in a couple of minutes while in cases where a network was also provided it only took a few seconds. The two most time consuming steps were inferring the network in cases one was not provided, and matching not-Silva, non-GTDB taxonomies to GTDB genomes. When microbetag’s preprocessing was used, and its results were provided as input, the computational time for the on-the-fly part was again only a couple of seconds even for larger networks. The computing requirements for running microbetag with custom genomes are strongly dependent on the number of bins/MAGs/genomes and whether annotation steps and GEM reconstruction have been already performed or not.

Yet, there are several challenges involved in our approach. First, microbetag inherits all the biases and drawbacks of both the data and the software it is based on. Functional annotation comes with its own limitations. Some functional domains boast richer annotations and more comprehensive descriptions compared to others, thus exhibiting a wealth of detail and employing more precise terminology, particularly for widely recognized processes (e.g., glycolysis versus secondary metabolites biosynthesis).

In the validation case, the bin representing the *Variovorax* strain was mapped to a genome that is supposed to contain the pantothenate KEGG modules. Thus, the fact that it requires pantothenate to grow, as the authors mention, would not have been predicted in the microbetag framework. This highlights the importance of strain-level variety and also the difference between genomic potential versus expression and synthesis of enzymes. Beyond the sequencing and annotation challenges, we also need to consider the fact that a pathway may not be fully represented in a KEGG module or that an organism may have alternative pathways for its product(s).

Pathway complementarity can only be as accurate as the KEGG MODULE database and as precise as the software annotating genomes with KO terms. It is well known that automated metabolic network construction comes with a number of challenges and different software for this task have their own limitations (40). Using ModelSEED with a complete medium may limit potential metabolic interactions but the retrieved ones will be more reliable because using a complete medium to gapfill a metabolic network reduces the number of reactions that need to be added.

In addition, pathway complementarity *per se* does not guarantee that intra-cellular metabolites are indeed exchanged. microbetag does not check whether metabolites can be excreted or consumed. If the donor lacks a transporter and the metabolite cannot cross its cell wall, sharing could still occur through lysis, but that requires a sufficiently high death rate. Furthermore, the seed approach depends on an external definition of metabolite essentiality to define seeds, but it is not always known whether metabolites that cannot be synthesised without seeds are indeed essential to growth. These are all reasons why pathway and network complementarities are predictions of potential cross-feeding relationships and do not necessarily reflect actual interactions.

In addition, pairwise relationships do not capture higher-order ecological interactions, in which species depend on (or are influenced by) multiple other species (1). However, since microbetag is decoupled from network inference, it could annotate a network with hyperedges (i.e. edges connecting more than two taxa) produced by a future tool capable of inferring higher-order interactions.

Last, the limited number of Archaea in microbetagDB is also the result of a software limitation. As shown in (41) (see Fig. 6b), the original version of CheckM (42) that is still being used by GTDB returns lower completeness scores for genomes that correspond to phyla known for having smaller genomes in general, e.g. Patescibacteria representative genomes in GTDB have an average completeness of ∼65%. Thus, only a few representatives from these taxonomic groups passed our filters leading to an important under-representation of Archaea.

### Context

microbetag is among a number of tools that are based on the reverse-ecology approach, whose goal it is to derive ecological insight, in particular on interactions, from the genomes of community members (25,43–45). microbetag goes beyond these previous tools and methods by combining interaction prediction based on metabolic networks not only with microbial network inference but also with the systematic annotation of taxa with phenotypic properties. In addition, it makes all these approaches accessible to researchers without bioinformatics skills. As such, it is a unique resource of potential use to everyone working with microbiome data.

### Future work

In the near future, we plan to develop two main features: a) the integration of transcriptomics data provided by the user, which would enhance or lower the probability for a potential metabolic interaction to occur based on whether the KO terms involved are present or not, and b) the integration of spatial data; it is well-known that the distance between cells determines whether an interaction occurs (46). For this, we intend to support data with spatial dimensions. We may also consider the integration of additional phenotype prediction tools, such as bacLIFE (47).

## Conclusions

Co-occurrence networks are widely used in microbiome studies to explore associations. However, their inference and their interpretation come with several challenges. Metabolic exchanges among microbial taxa are considered ubiquitous (48) in a large number of environments. In our study, we exploit reverse-ecology approaches and publicly available genomic data and software to predict phenotypic traits and construct metabolic networks to annotate co-occurrence networks derived from amplicon or shotgun data. Our annotation was in agreement with the study of Hessler et *al.* (32) predicting thiamine-related metabolic interactions among *Variovorax* and its closest neighbors, suggesting several ways to achieve them. Using the Cabrera et *al.* Dataset (38), we showcased the potential of microbetag as a hypothesis-generating tool suggesting mechanisms for the increase of butyrate producers in infant microbiota supplemented with iron and GOSFOS. microbetag is a first one-stop-shop platform for the annotation of microbial co-occurrence networks, highlighting the potential of data integration combined with follow-up analyses for network interpretation.

## Methods

### Genomes included

Using the Genome Taxonomy Database (GTDB) v207 metadata files, we retrieved the NCBI genome accessions of the high-quality representative genomes, i.e., completeness ≥95% and contamination ≤5%. A set of 26,778 genomes was obtained, representing 22,009 unique NCBI Taxonomy Ids. Using these accession numbers, we were able to download their corresponding .faa files when available, leading to a set of 16,900 amino acid sequence files. The latter were annotated (see “Genome-based node annotation”) and used to obtain potential pathway complementarities between pairs of genomes (see “Pathway Complementarity”). Last, when available, their corresponding annotations in the PATRIC database (49) were retrieved to reconstruct metabolic networks (see “Seed scores and complements”).

### Taxonomy schemes

microbetag maps the taxonomy of each entry in the abundance table to its corresponding NCBI Taxonomy Id and, if available, its closest GTDB representative genome(s). Several GTDB representative genomes may map to the same NCBI Taxonomy Id. Two well established taxonomy schemes are supported: the GTDB (50) that is being broadly used for bins and/or MAGs (Metagenome Assembled Genomes) taxonomic classification and the Silva database (51) that is widely used in amplicon studies. Both taxonomy schemes link their taxonomies to NCBI Taxonomy Ids (52). In case neither of those two taxonomies was used, and the abundance table contains less than 1,000 taxa, microbetag maps the user-provided taxonomies to the NCBI Taxonomy. To this end, microbetag makes use of the fuzzywuzzy library (v. 0.18.0) that implements the Levenshtein Distance Metric to get the closest NCBI taxon name and thus its corresponding NCBI Taxonomy Id; a high similarity score is used [90] to avoid false positives. In the rare case a taxon receives the same score with more than one NCBI taxonomy, microbetag will choose one at random. Also, using the nodes dump file of NCBI Taxonomy, microbetag may retrieve the child taxa of a taxon in user data, along with their corresponding NCBI Taxonomy Ids, if requested by the user. If the user provides their abundance table with taxonomies already mapped to the GTDB taxonomy, microbetag will quickly report the best possible annotations.

### Network inference

When a microbial network is not provided by the user, microbetag relies on FlashWeave (v. 0.19.2) (8) to build one on the fly. microbetag supports the annotation of networks built from any algorithm/software, in any format Cytoscape can load.

### Microbetag preprocessing

To aid the user to map their sequences to the GTDB taxonomy, DADA2-formatted 16S rRNA gene sequences for both bacteria and archaea (53) were used to train the IDTAXA classifier of the DECIPHER package (v. 2.14.0) (54) and are available through the microbetag preprocess Docker image. Likewise, when the abundance table consists of more than 1,000 taxa and/or the taxonomy scheme is not among the automatically mapped, providing a network as an input is mandatory. The microbetag preprocess Docker image also supports the inference of a network using FlashWeave.

### Literature-based node annotation

Using a set of Tara Ocean samples (55), FAPROTAX (56) estimates the functional potential of the bacterial and archaeal communities, by classifying each taxonomic unit into functional group(s) based on the current literature, descriptions of cultured representatives and/or manuals of systematic microbiology. In this manually curated approach, a taxon is associated with a function only if all the cultured species within the taxon have been shown to exhibit that function. In the version microbetag makes use of, FAPROTAX (v. 1.2.4) includes more than 80 functions based on 7600 functional annotations and covers more than 4600 taxa. Contrary to gene-content-based approaches, e.g., PICRUSt2 (57), FAPROTAX estimates metabolic phenotypes based on experimental evidence. microbetag invokes the accompanying script of FAPROTAX and converts the taxonomic microbial community profile of the samples included in the user’s abundance table or of the taxa present in the provided network, into putative functional profiles. Then, it parses FAPROTAX’s sub-tables to annotate each taxonomic unit present in the user’s data with all the functions for which they had a hit. FAPROTAX annotations are not part of the microbetagDB but are computed on the fly.

### Genome-based node annotation

phenDB (58) is a publicly available resource that supports the analysis of bacterial (meta)genomes to identify 47 distinct functional traits, e.g. whether a species is producing butanol or has a halophilic lifestyle. It relies on support vector machines (SVM) trained with manually curated datasets based on gene presence/absence patterns for trait prediction. More specifically, the model for a particular trait is trained using a collection of EggNOG annotated genomes (59) where the knowledge of whether that trait is present or absent among its members is available. These models (classifiers) are used to predict presence/absence of their corresponding traits in non-studied species.

For microbetagDB, classifiers were re-trained using the genomes provided by phenDB for each trait to sync with the latest version of EggNOG (v. 5.0.0) (59) and the phenotrex (v. 0.6.0) (58) software tool. Genomes were downloaded from NCBI using the Batch Entrez program. Then, *genotype* files were produced for all the high quality GTDB representative genomes. Each model was then used against all the GTDB *genotype* files to annotate each with the presence or the absence of the trait. A list of all the phenotypic traits available for the genomes present in microbetagDB is available on microbetag’s documentation site. The updated models are also available. Figure 5 summarizes these precalculation steps.

**Fig. 5:**
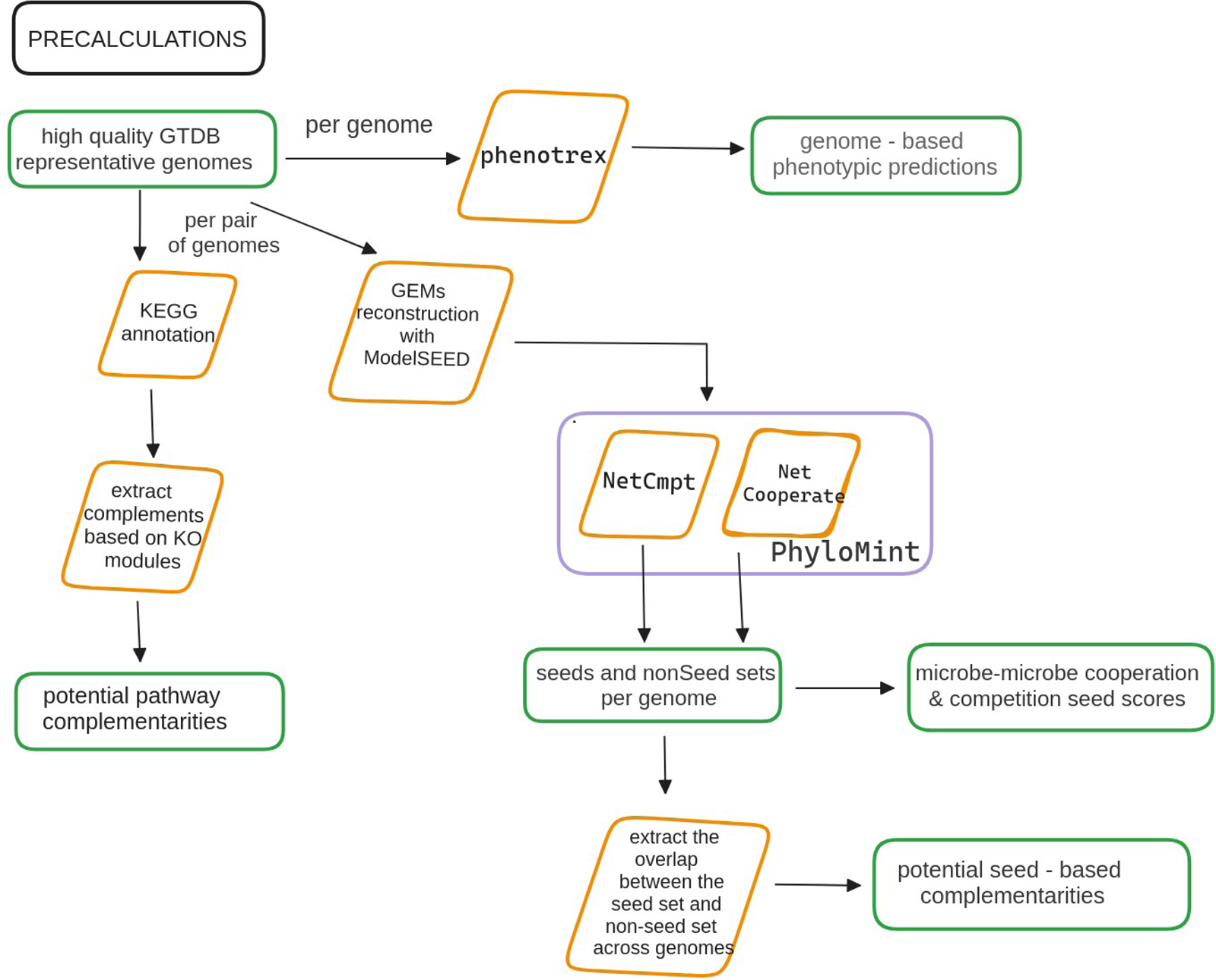
Diagram of the microbetag precalculations. GTDB v207 representative genomes were filtered and for those of high quality, 33 phenotypic traits were predicted using phenotrex. To this end, models were re-trained to sync with version 5.0 of EggNOG.

### Pathway complementarity

To infer potential pathway complementarities, we rely on the modules in the KEGG MODULES database (27). A KEGG module is defined as a functional unit within the KEGG framework that represents a set of enzymes and reactions involved in a specific biological process or pathway (27). Such a unit consists of several *steps*, each of which may have more than one molecular way to occur (Fig. 6). A module’s definition is a logical expression and consists of KEGG ORTHOLOGY terms (KOs) that may be coupled with one another as: a) connected steps of the pathway, b) parts of a molecular complex, c) alternatives of the same step, and d) optional entities of a complex. Both (a) and (b) cases should be considered as the AND logical operator, while (c) would be the OR (Fig. 6).

Given a module’s definition, we will consider as an *alternative* any subset of the KO terms mentioned in the definition, that has exactly one way to perform each step, provided that all the steps of the module are covered. We define a genome as having a *complete* module, only if all the KOs of at least one alternative are present in it. In Supplementary Information, we show an example of a module along with its alternatives. Within this framework, kofamscan (v. 1.3.0) (60) was used to annotate with KOs the 16,900 high quality GTDB representative genomes for which a .faa file was available (61). The KOs of each genome were then mapped to their corresponding KEGG modules; a KO may map to more than one module (1:*n*).

All module definitions were retrieved using the KEGG API and parsed to enumerate their alternatives. Each pair of the KEGG annotated genomes was then investigated for potential pathway complementarities, i.e., whether a genome lacking a number of KOs (*genome_A_*) to have a complete module (*module_x_*) could benefit from another species’ genome(s) (*genome_B_*). In that case, *genome_B_* does not necessarily have a complete alternative of *module_x_*; as long as it has the missing KOs that *genome_A_* needs to complete an alternative of it, *genome_B_* potentially complements *genome_A_* with respect to *module_x_*.

Thanks to the graphical user interface (GUI) of the KEGG pathway map viewer (62,63), each complementarity can be visualized as part of a KEGG metabolic map. In this case, the KOs contributed by the donor are shown in blue-green whereas those coming from the beneficiary genome are colored in red.

microbetag annotates the edges of a microbial network by identifying pairs where both taxa map to an annotated genome present in microbetagDB. Since microbial networks are usually undirected, both nodes of an association are considered as potential donors and beneficiary species. When more than one GTDB representative genome maps to the same NCBI Taxonomy Id all the possible genome combinations are considered. Finally, two edges are added for such taxon pairs in the annotated network: one considering *species_A_* as the potential beneficiary and *species_B_* as the potential donor species, and one vice-versa.

**Fig. 6:**
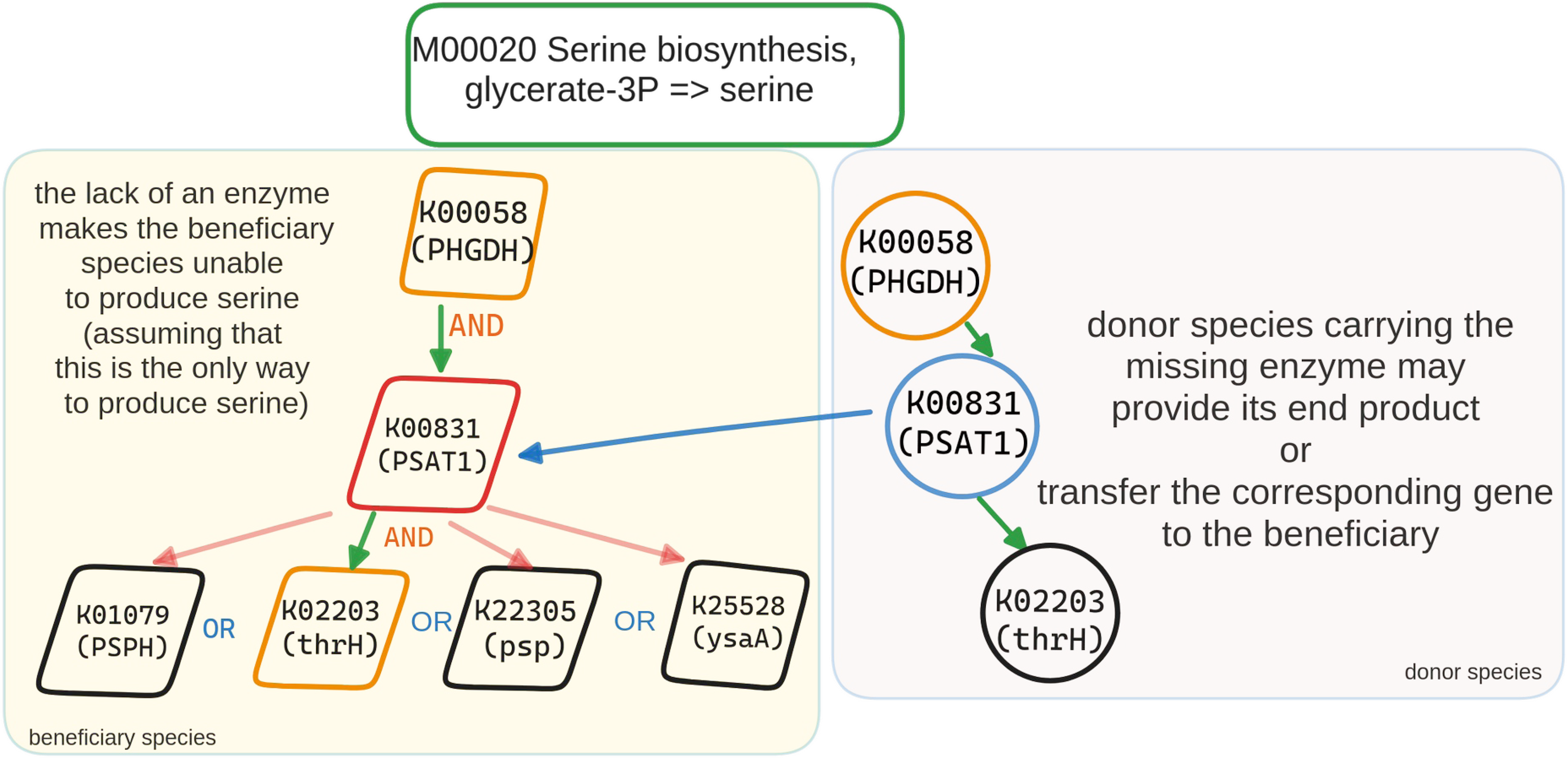
Pathway complementarity approach. The high-quality GTDB genomes were annotated with KEGG ORTHOLOGY (KO) terms. All the possible ways a donor species could “fill” a beneficiary’s incomplete KEGG module were calculated. In this case, there are four unique ways to complete the serine biosynthesis module; in all of them K00831 is required, which is missing from the beneficiary species that supports two out of the three steps of the module’s definition. The enzyme linked to KOO831 in the donor species completes the glycerate-3P to serine biosynthesis pathway in the beneficiary. Such complementarities may represent interactions in case the beneficiary depends on the end product to grow, has no alternative pathways to synthesize it and intermediary metabolites can be shared.

### Seed scores and complements using genome scale metabolic reconstructions

The Metabolic Complementarity Index (*MI_Complementarity_*) measures the degree to which two microbial species can mutually assist each other by complementing each other’s biosynthetic capabilities. As described in (64), it is defined as the proportion of seed compounds of a species that can be synthesized by the metabolic network of another species but are not included in the seed set of the latter. *MI_Complementarity_* offers an upper bound assessment of the potential for cross-feeding interactions between two species. Further, the Metabolic Competition Index (*MI_Competition_*) represents the similarity in two species’ nutritional profiles. This index establishes an upper limit on the level of competition that one species may face from another. Those indices have been described and implemented in the NetCooperate (24) and NetCmpt (23) tools, respectively. We will be referring to those two indices as “seed scores”. Recently, the PhyloMint tool (v. 0.1.0) (64) was released supporting the calculation of the seed scores of metabolic networks in SBML format.

In the microbetag framework, seed scores were computed using metabolic networks derived from the high-quality GTDB representative genomes and the PhyloMint tool. Metabolic networks were reconstructed using the Model SEED pipeline (65) through its Python interface <u>ModelSEEDpy</u> (v. 0.2.2). The latter requires RAST annotated genomes (66); if available through the PATRIC database (49), annotations were retrieved. For the rest of the genomes, RAST annotation was performed through RASTtk (v. 1.3.0) (67).

In contrast to seeds, non-seeds are metabolites that the metabolic network can produce on its own. The seed and non-seed sets of each genome were used to compute their overlap among all the pairwise combinations of those genomes. More specifically, seed and non-seed compounds of each metabolic network were mapped to their corresponding KO terms (as substrates or products of their reactions) and those related to a KEGG MODULE were considered further. Focusing on the KEGG

MODULE-related KO terms as terms of interest, the overlap of *seed set_speciesA_* with the *non seed set_speciesB_* was retrieved (Fig. 7). Such *seed complementarities* were calculated for all metabolic network pairs and are now available through microbetagDB. Edges of microbial networks where both taxa have been mapped to at least one GTDB genome are further annotated by listing all the KEGG maps for which there is at least one seed compound of the potentially beneficiary species.

**Fig. 7.**
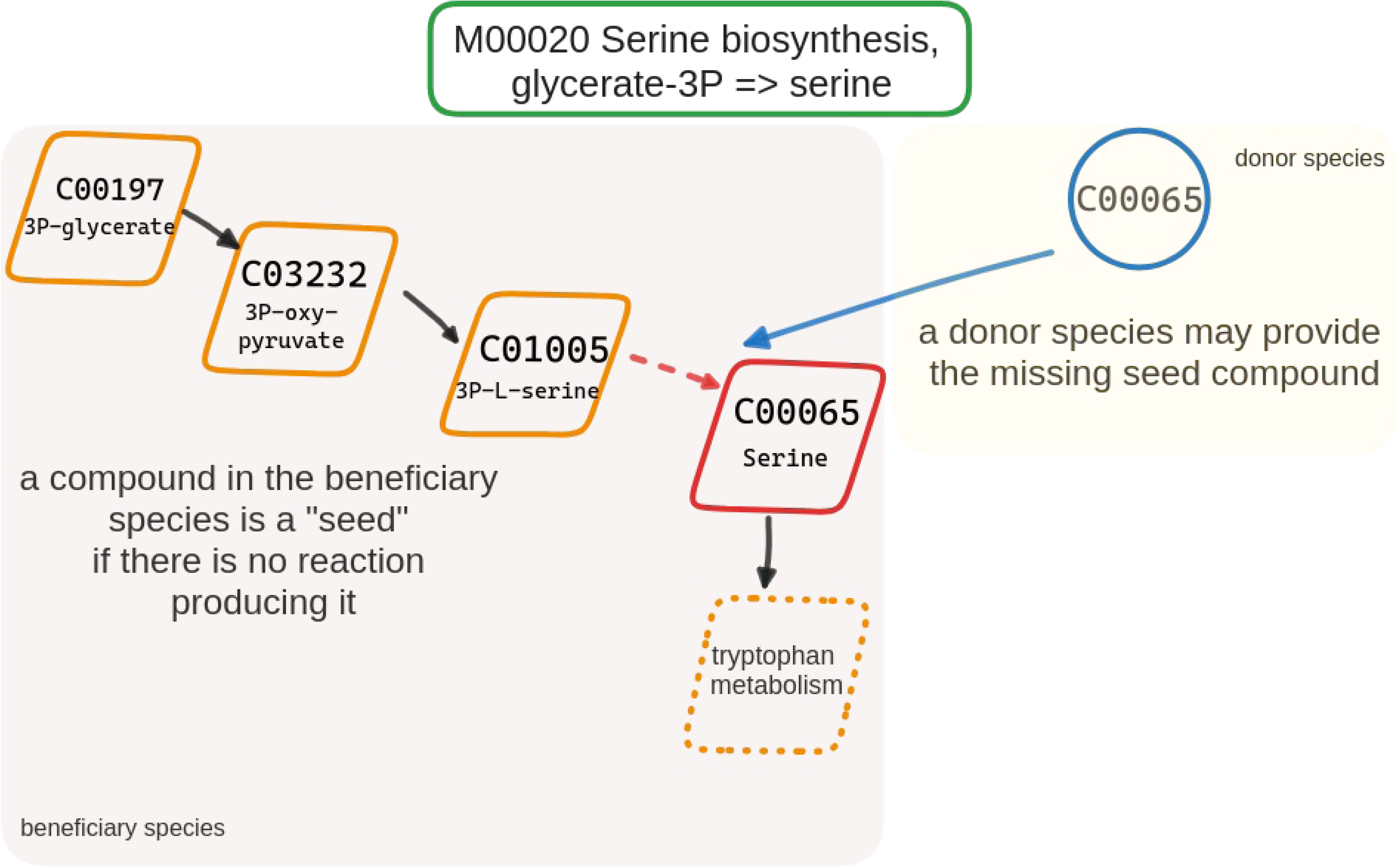
Seed complementarity. Using the entire metabolic network of a species, compounds can be identified that are required for growth but cannot be produced by the species’ metabolism (“seeds”). Such compounds are taken up from the environment and could be provided through cross-feeding. In the case shown here, species A requires serine but cannot produce it on its own. Thus, a potential donor species B may provide it.

### Network clustering

manta (15) is a heuristic network clustering algorithm that clusters nodes within weighted networks effectively, leveraging the presence of negative edges and discerning between weak and strong cluster assignments. microbetag invokes manta (<u>edited version</u> of v. 1.1.1) to cluster the microbial network. In case manta was executed, the annotated network inherits the layout that manta returns. However, as mentioned below, the microbetag GUI (MGG) can work with clusters resulting from any network clustering algorithm if it meets the expected output format.

### The microbetag workflow

As shown in Fig. 8, the microbetag workflow expects an abundance table representing either amplicon or shotgun data. If a microbial network is already available, the user may provide it too as input. The microbetag workflow will first map the taxa present in the abundance table to their corresponding GTDB representative genomes if that is possible, i.e., in case the taxonomy provided does reach the species or the strain level (see “Taxonomy schemes”). If a network is not provided, microbetag will build one using FlashWeave. Then the abundance table will be used for a literature-based annotation using FAPROTAX. This is the only annotation step that is microbetagDB-independent within the web-service workflow. The nodes of the network will be further annotated with phenotypic traits based on phenDB predictions (58). Edges linking taxa assigned to the species or strain level will be annotated with pathway and seed complementarities and seed scores. Last, network clustering will be performed with manta, assigning each node to a cluster. The annotated network is then returned in a .cx2 format. The user may skip any of these annotation steps if not needed for their analysis.

**Fig. 8.**
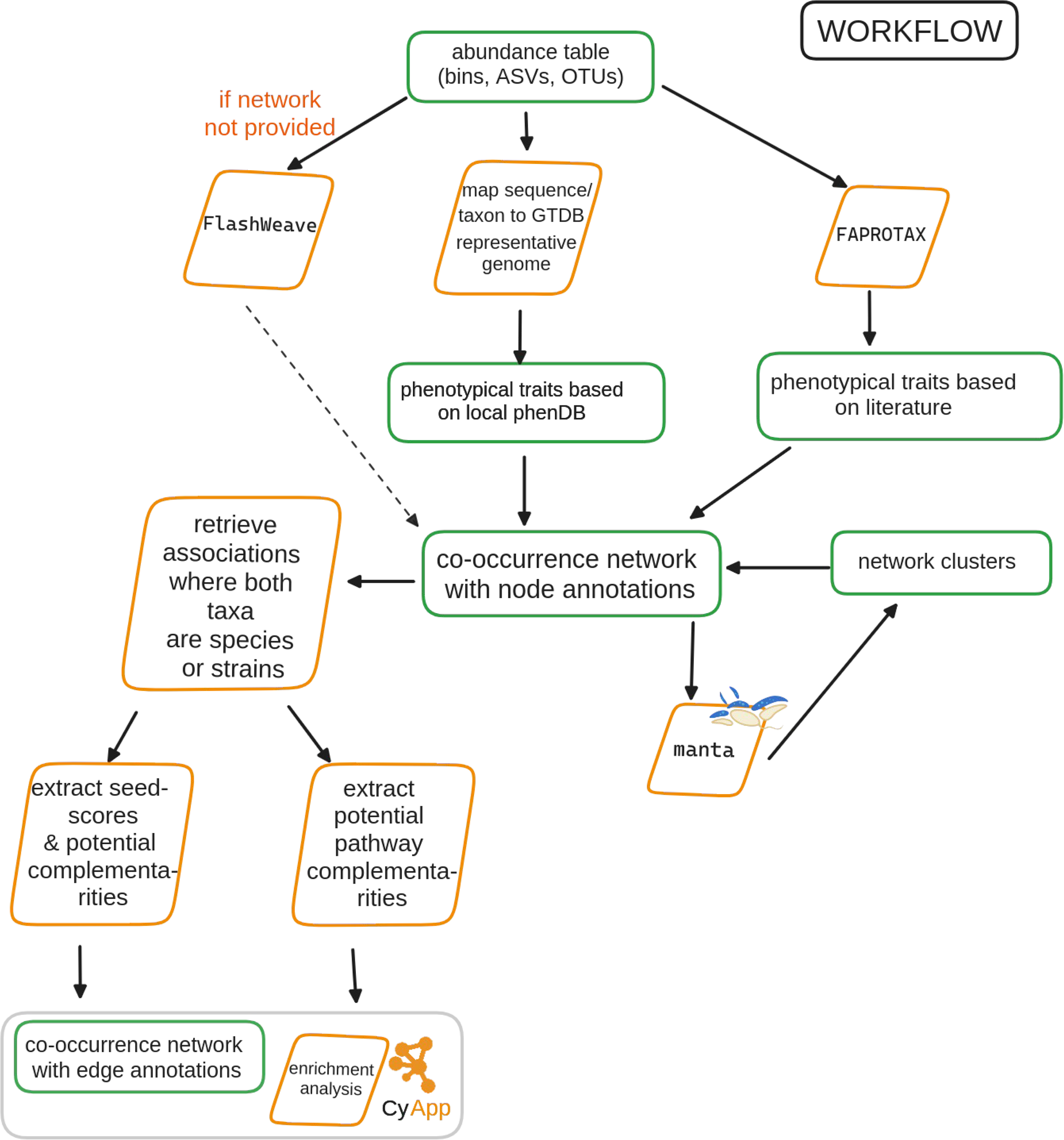
Diagram of microbetag’s on-the-fly workflow. microbetag expects either an abundance table only as input and infers a microbial network using FlashWeave or an abundance table along with an already inferred network. After mapping taxa to GTDB reference genomes, for those with sufficient taxonomic resolution, phenotypic attributes are assigned to the nodes. Literature-based annotations of the nodes are also added using FAPROTAX. On the edge level, microbetag assigns the precalculated potential complements based on the pathway and the seed complementarity approaches. microbetag supports optional network clustering with manta. The annotated network can then be parsed into Cytoscape using the MGG app.

### The MGG Cytoscape App

To enable a straightforward, user-friendly way to perform the microbetag workflow and visualize microbetag-annotated networks we developed a Cytoscape app. The microbetag GUI (MGG) app was built based on the source code of the scNetViz (68) (forked from commit <u>336ac63</u>). A visual style was developed to better distinguish annotated nodes and edges. Nodes are colored based on the level of the taxonomic assignment with those being annotated highlighted in green. Similarly, edges are green when they carry a positive weight and dark pink when negative. Black edges denote pathway and/or seed complementarities. The latter were not accounted for in the edge weights since edges represent an undirected relationship while complementarity/overlap scores assume a direction, i.e. the complementarity score of species A versus B is not necessarily the same as that of B versus A.

### Groups of phenotypic traits

Phenotypic traits returned from FAPROTAX and phenDB-like annotation steps are organized into biologically meaningful groups. The main groups supported are related to a) the lifestyle of a species, for example being halophilic or thermophilic etc., b) the biogeochemical processes that are linked to the metabolic potential of a species, for example Nitrite-oxidizing bacteria (NOB) bacteria and c) important metabolites a species is predicted to produce, e.g. butanol. The aim of these groups is to facilitate filtering of the taxa present. Enrichment analysis for traits in such groups (e.g., based on the clusters identified by an algorithm like manta) can be performed through the Cytoscape app.

### Software architecture

microbetag is a Docker-based application. We deployed the microbetag application using Docker containers (69) (v24.0.2) managed by Docker Compose (see Supplementary Fig. 1). Docker Compose is a tool for defining and running multi-container Docker applications using a YAML file to configure the services required for the application. Containers of three Docker images are being used simultaneously: a) a MySQL database including the microbetagDB, b) a nginx (70) web server and c) the application itself, including the API and the microbetag workflow. Gunicorn (20.1.0) was used to build an application server which communicates with the web server using the Web Server Gateway Interface (WSGI) protocol and handles incoming HTTP requests. microbetag is implemented as a Flask application (v2.3.2); Flask is a micro web framework for developing Python web applications and RESTful APIs. The API has a route for performing the microbetag workflow, either through any Python console or the Cytoscape MGG app, but also several other routes that enable quick and easy access to the microbetagDB content, i.e. the genomes present, their phenotypic traits predicted based on genome annotations, pathway and seed complementarities among specific genomes or NCBI Taxonomy Ids and their corresponding seed scores if available. A thorough description of the microbetag API is available at the documentation web site. The source code of the microbetag web service is available on GitHub (see Code availability).

## Declarations

Availability of data and materials

– Raw sequences for the use case: The sequencing datasets generated by Cabrera et al. are available in the European Nucleotide Archive (ENA) repositories

PRJEB29350 (https://www.ncbi.nlm.nih.gov/bioproject/513540),

PRJNA279279 (https://www.ncbi.nlm.nih.gov/bioproject/PRJNA279279) and PRJNA629336 (https://www.ncbi.nlm.nih.gov/bioproject/PRJNA629336)

– Raw data for the validations case: The sequencing datasets generated by Hessler et al. are available in the European Nucleotide Archive (ENA) repository (https://www.ebi.ac.uk/ena/browser/view/PRJEB67393)

– All files for the tutorials and the computing times mentioned can be found at microbetag’s GitHub repository: https://github.com/hariszaf/microbetag/tree/gh-pages/download

## Funding

This work was initiated thanks to an EMBO Scientific Exchange Grant to HZ. It was then supported by the 3D’omics Horizon 2020 project (101000309). We would also like to thank the National Resource for Network Biology (NRNB) and the Google Summer of Code 2023 for the support of E.I.M.D.

## Conflict of interest/Competing interests

The authors declare that they have no other competing interests.

## Authors’ contributions

Conceptualization: K.F. Methodology: K.F. and H.Z. Software: H.Z., E.I.M.D. and J.M. Validation: H.Z., K.F and A.S. Formal analysis: H.Z. and K.F. Investigation: H.Z., A.S. Resources: K.F., A.E. and A.G. Data Curation: H.Z. Writing - Original Draft: H.Z. and K.F. Writing - Review & Editing: all Visualization: H.Z. Supervision: K.F., H.Z. and S.M. Project administration: K.F. Funding acquisition: K.F., H.Z.

## Acknowledgments

We would like to thank Dr. Christina Pavloudi for the insight on how to organize the trait groups. We would also like to thank Dr. Hessler and Prof. Jillian F. Banfield for sharing both the bins and the network of their study.

## Ethics approval

Not applicable

## Consent to participate

Not applicable.

## Code availability

– microbetagDB related scripts: https://github.com/hariszaf/microbetag
– microbetag application: https://github.com/msysbio/microbetagApp
– MGG CytoscapeApp: https://github.com/ermismd/MGG/
– Validation and use case: https://github.com/hariszaf/microbetag/tree/ms/docs
– Documentation web site: https://hariszaf.github.io/microbetag/

## Notes

https://hariszaf.github.io/microbetag/

https://github.com/hariszaf/microbetag

https://doi.org/10.5281/zenodo.10562677

https://apps.cytoscape.org/apps/mgg

### Competing Interest Statement

The authors have declared no competing interest.

