## Supplementary Information for "microbetag: simplifying microbial network interpretation through annotation, enrichment tests and metabolic complementarity analysis"

### Background on pathway and seed complementarity

For a genome to have a KEGG module *complete* means it provides at least one complete *alternative*. Alternatives are considered as the unique combinations of KOs that connect an input compound to an output compound of the module

For example, the definition of the D-Galacturonate degradation in Bacteria ([M00631)](https://www.genome.jp/dbget-bin/www_bget?M00631) is:

K01812 K00041 (K01685,K16849+K16850) K00874 (K01625,K17463)

Once breaking down, it leads to 4 alternative sets of KOs (pathways):

## K01812 K00041 K01685 K00874 K01625 K01812 K00041 K16849+K16850 K00874 K01625 K01812 K00041 K01685 K00874 K17463 K01812 K00041 K16849+K16850 K00874 K17463

In alternatives two and four, the K16849+K16850 is a *complex*, meaning both KO terms are required for the step to be available.

In case of seed complementarity, in microbetag we focus on the effect that a metabolic exchange between two taxa might have if the seed of the beneficiary taxon is linked to a KEGG MODULE. Therefore, the KOs that were found linked to modules were mapped to ModelSEED ids. The initial seed and non-seed sets that were exported as sets of ModelSEED ids were then mapped to KOs too. When the non-seed set of a genome (donor) provided a seed related to a KEGG module to another genome (beneficiary), this is considered a potential metabolic interaction.

**Supplementary Table 1:** Bin sequence files were mapped against GTDB using GTDB-tk. Chloroflexi refers to the GTDB taxonomy of: 54-19 sp001898225

| **Potential thiamine-related complements among Variovorax neighbors** | | |
| --- | --- | --- |
| **Beneficiary** | **Donor** | **potential complement** |
| *Terrimonas ferruginea* | *Tahibacter* sp001725155 | [C01081](https://www.genome.jp/dbget-bin/www_bget?cpd:C01081) |
| *Terrimonas ferruginea* | *Rhodoglobus* sp001725325 | [C01081](https://www.genome.jp/dbget-bin/www_bget?cpd:C01081) |
| *Nitrosospira* sp001899235 | *Bosea* sp001898115 | [C04327](https://www.genome.jp/dbget-bin/www_bget?cpd:C04327);[C01279](https://www.genome.jp/dbget-bin/www_bget?cpd:C01279) |
| Chloroflexi | *Bosea* sp001898115 | [C15809](https://www.genome.jp/dbget-bin/www_bget?cpd:C15809) |
| Chloroflexi | Xanthobacteraceae | [C15809](https://www.genome.jp/dbget-bin/www_bget?cpd:C15809) |
| Chloroflexi | *Nitrosospira* sp001899235 | [C15809](https://www.genome.jp/dbget-bin/www_bget?cpd:C15809) |

**Supplementary Table 2**: *Variovorax* genomes present in microbetagDB and their corresponding complete/incomplete presence of the pantothenate related KEGG modules. The genome that bin_55 was mapped to is shown in bold.

| **Genome** | **md:M00119** | **md:M00913** |
| --- | --- | --- |
| GCA_004210915.1 | incomplete | complete |
| GCA_902506565.1 | incomplete | incomplete |
| GCF_000184745.1 | complete | complete |
| GCF_000282635.1 | complete | complete |
| GCF_000463015.1 | complete | complete |
| GCF_000834655.1 | complete | complete |
| GCF_001424835.1 | complete | complete |
| GCF_001425205.1 | complete | complete |
| GCF_001426505.1 | complete | complete |
| GCF _001577265.1 | incomplete | incomplete |
| GCF _002157355.1 | complete | complete |
| GCF_002754375.1 | complete | complete |
| GCF_003019815.1 | incomplete | complete |
| **GCA_001899795.1** | **complete** | **complete** |
| GCF_003852515.1 | complete | complete |
| GCF_003951285.1 | complete | complete |
| GCF _003952165.1 | complete | complete |
| GCF_003952185.1 | complete | complete |
| GCF_003984625.1 | complete | complete |
| GCF_003984645.1 | complete | complete |
| GCF_006438845.1 | complete | complete |
| GCF_007828835.1 | complete | complete |
| GCF_009498455.1 | complete | complete |
| GCF_009755665.1 | complete | complete |
| GCF_010499245.1 | complete | complete |
| GCF_013376045.1 | complete | complete |
| GCF_014170375.1 | complete | complete |
| GCF_014302995.1 | complete | complete |
| GCF_014303735.1 | incomplete | incomplete |
| GCF_901827175.1 | complete | complete |
| GCF_ 901827205.1 | complete | complete |

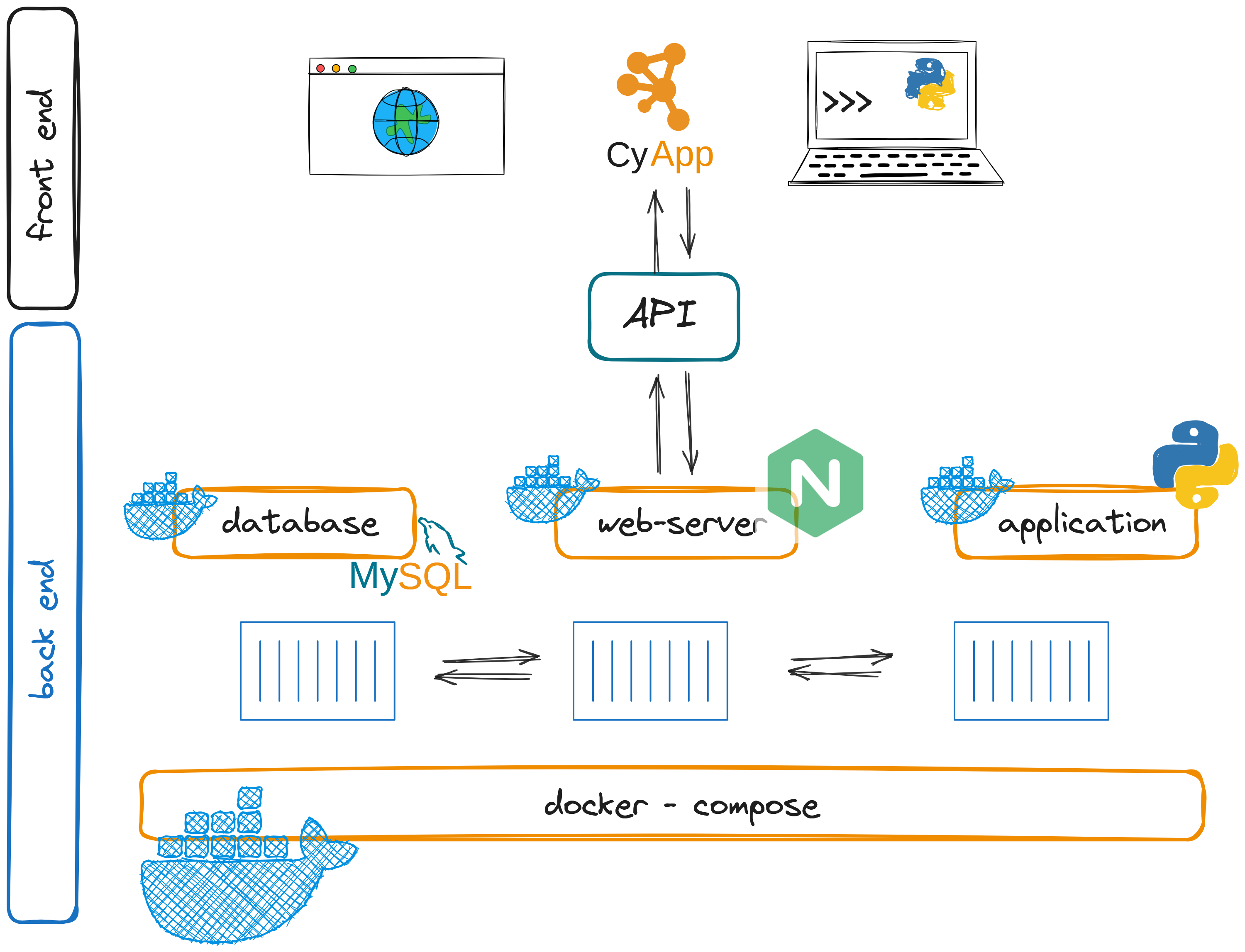

**Supplementary Fig. 1**: microbetag software ecosystem. Three Docker containers are combined: a nginx web server connected to a MySQL database and the microbetag workflow. An API, as part of the last container, enables communication between the client (front) and the server side (back end). The content of the microbetagDB and the microbetag workflow are accessible through a web-browser, a terminal and a CytoscapeApp (MGG).

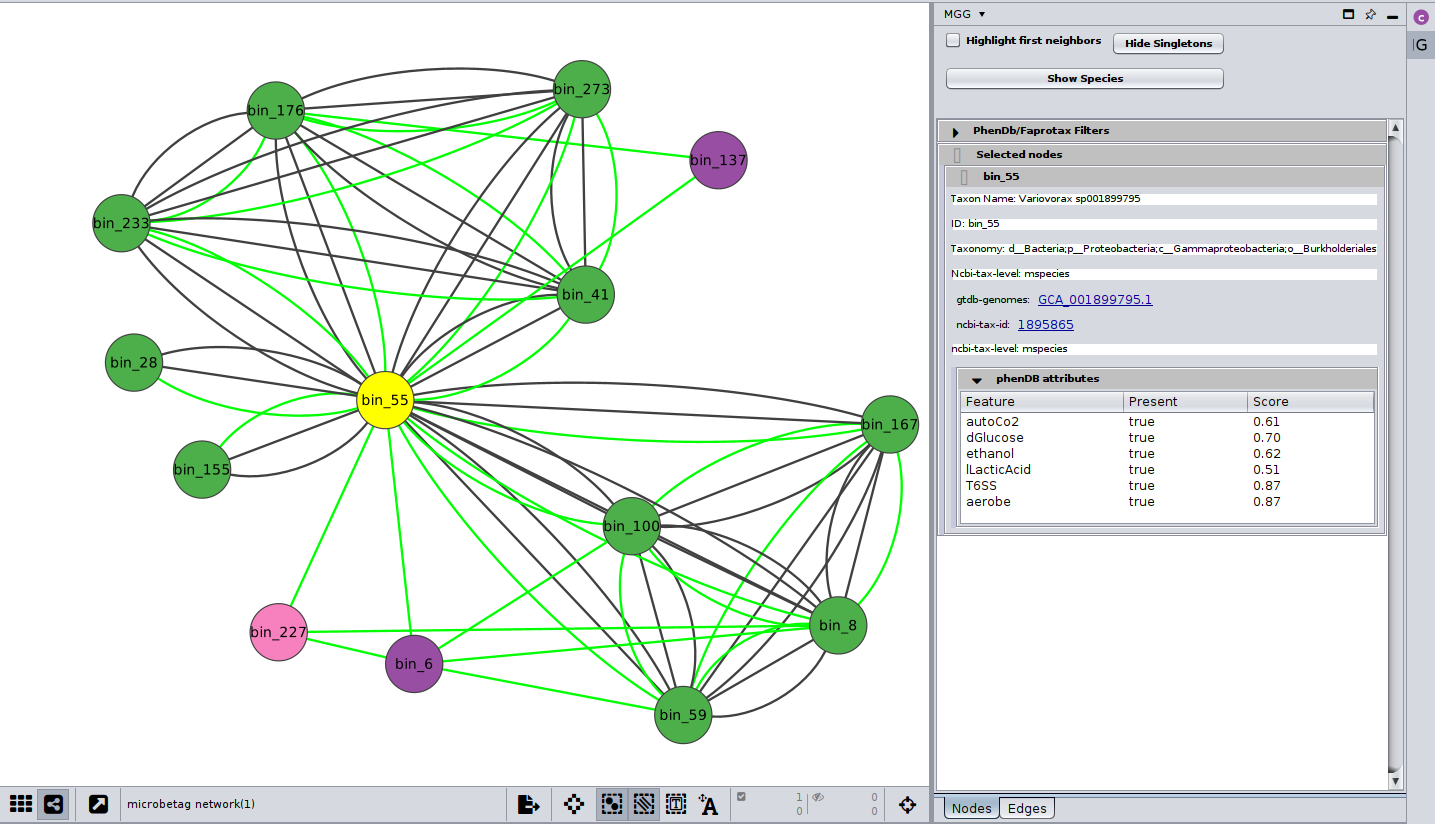

**Supplementary Fig. 2**: *Variovorax* node (bin_55) and its neighbors annotated by microbetag. Only three of them were not mapped to a GTDB representative genome (pink and purple nodes denoting genus and family taxonomic levels accordingly). Green edges represent the positive association weights. The black edges represent pairwise seed complementarities and scores.
